# Priority-based transformations of stimulus representation in visual working memory

**DOI:** 10.1101/2021.05.13.443973

**Authors:** Quan Wan, Jorge A. Menendez, Bradley R. Postle

## Abstract

How does the brain prioritize among the contents of working memory (WM) to appropriately guide behavior? Using inverted encoding modeling (IEM), previous work (Wan et al., 2020) showed that unprioritized memory items (UMI) are actively represented in the brain but in a “flipped”, or opposite, format compared to prioritized memory items (PMI). To gain insight into the mechanisms underlying the UMI-to-PMI representational transformation, we trained recurrent neural networks (RNNs) with an LSTM (long short-term memory) architecture to perform a 2-back working memory task. Although visualization of LSTM hidden layer activity using Principal Component Analysis (PCA) suggested that stimulus representations undergo a smooth rotational transformation across the trial, demixed (d)PCA of the same data decomposed this pattern into a cascade of multiple trajectories, each with a different time course, unfolding within UMI and PMI subspaces. The application of the same analyses to the EEG dataset of Wan et al. (2020) indicated that an item’s trajectory through the UMI subspace closely mirrored that of the RNN, but that its trajectory through the PMI subspace differed markedly from the RNN. It may be a general principle that, at the level of the representational code, information held in WM undergoes priority-based transformations that allow for its retention while preventing it from interfering with concurrent behavior. Implementational details of this process may vary across model systems.

**Author Summary:** How is information held in working memory (WM) but outside the current focus of attention? Motivated by previous neuroimaging studies, we trained recurrent neural networks (RNNs) to perform a 2-back WM task that entails shifts of an item’s priority status. Dimensionality reduction of the resultant activity in the hidden layer of the RNN allowed us to characterize how a stimulus item’s representation follows a transformational trajectory through high-dimensional representational space as its priority status changes from memory probe to unprioritized to prioritized. This work illustrates the value of artificial neural networks for assessing and refining hypotheses about mechanisms for information processing in the brain.

## Introduction

The ability to flexibly select and prioritize among information held in working memory (WM) is critical for guiding behavior and thought. For this reason, the neural mechanisms of attentional prioritization in WM have been extensively studied in recent years. Many studies of WM for visual material have reported that the prioritization of one item held in WM leads to a decrease in the activity level of the “unprioritized memory item” (UMI; Myers et al., 2017), sometimes to baseline levels (LaRocque et al., 2012; Lewis-Peacock et al., 2011; Rose et al., 2016). Some have interpreted these results as consistent with the idea that whereas prioritized memory items (PMI) are held in an active state, UMIs may be maintained as “activity-silent” traces encoded in synaptic weights (Barak & Tsodyks, 2014; Stokes, 2015). Although this possibility remains a topic of vigorous debate (Christophel et al., 2018; Schneegans & Bays, 2017; Sprague et al., 2016; Stokes et al., 2020), the present report does not relate directly to this question. Instead, of primary relevance here are more recent empirical results suggesting that, rather than producing a decline to baseline, deprioritization may produce a representational transformation of an item into a different, but still active, format. (We will return to a consideration of activity-silent models in the Discussion.)

Experimental tasks used to study prioritization in WM necessarily include multiple steps, such that the information not needed for the impending response (i.e., the UMI) might nevertheless be needed to guide a subsequent response. This is often done with retrocues. For example, van Loon and colleagues (2018) acquired functional magnetic resonance imaging (fMRI) data while first presenting subjects two target images sequentially (e.g., first a flower then a cow), then indicating with a cue whether memory for the first or second presented image would be tested first. Had the cue been a “1”, subjects would next see a test array of six flowers and indicate whether the target flower appeared in the test array, and finally a test array of six cows. On this trial, the target cow spent time as UMI, because the cue indicated that memory for the flower would be tested first. When van Loon et al. (2018) applied multivariate pattern analysis (MVPA) to fMRI data from posterior ventral temporal lobe, they found that a decoder trained on trials when an item was a PMI performed statistically below chance when that item was a UMI. Furthermore, a representational dissimilarity analysis indicated that, within their set of 12 stimuli (four cows, four skates, four dressers), each item’s high-dimensional representation in one state (e.g., as a PMI) was maximally different from its representation in the other state (i.e., as a UMI). Using a similar retrocuing procedure, Yu, Teng and Postle (2020) found, with multivariate inverted encoding modeling (IEM) of fMRI data from early visual cortex, that the reconstructed orientation of a grating “flipped” when it was a UMI relative to a PMI (e.g., a 30° orientation reconstructed as 120° while a UMI). Furthermore, for data from the intraparietal sulcus (IPS), they observed that the IEM reconstruction of the location where an item had been presented also flipped when an item’s priority status transitioned to UMI.

Shifts of priority are also characteristic of continuous-performance tasks, for which shifts of priority are dictated by task rules rather than by explicit cues. One example, which provided the impetus for the work presented here, is the 2-back WM task from Wan and colleagues (2020; Figure 1). Electroencephalography (EEG) signals were recorded while subjects viewed the serial presentation of oriented gratings and judged for each one whether it was a match or a non-match to the item that had appeared two positions previously in the series. This task entails a predictable transition through priority states for each item: When an item *n* is initially presented, it serves as probe to compare against the memory of item *n – 2*; after the *n-*to-*n – 2* decision is made, item *n* becomes a UMI while item *n – 1* is prioritized for the upcoming comparison with *n + 1*. Next, once the *n + 1*-to-*n – 1* comparison is completed, item *n* becomes a PMI for its impending comparison with item *n + 2*. To analyze the EEG data, we trained an IEM on the raw EEG voltages from a separate 1-item delayed-recognition task, and tested it on the delay periods separating *n* and *n + 1* and separating *n + 1* and *n + 2* (i.e., when item *n* assumed the status of UMI, then PMI). The results, reminiscent of van Loon et al. (2018) and Yu, Teng and Postle (2020), indicated that the IEM reconstruction of the UMI was “flipped” relative to the training data, then transformed again when its status transitioned to PMI (Figure 2). We referred to the transition from PMI to UMI, and back, as “priority-based remapping” (rather than “recoding” or “code morphing”; c.f. Parthasarathy et al., 2017), reasoning that the IEM reconstruction of the UMI would fail if it were represented in a neural code different from the trained model. To gain mechanistic insight into this phenomenon, formal modeling is needed.

**Figure 1.**
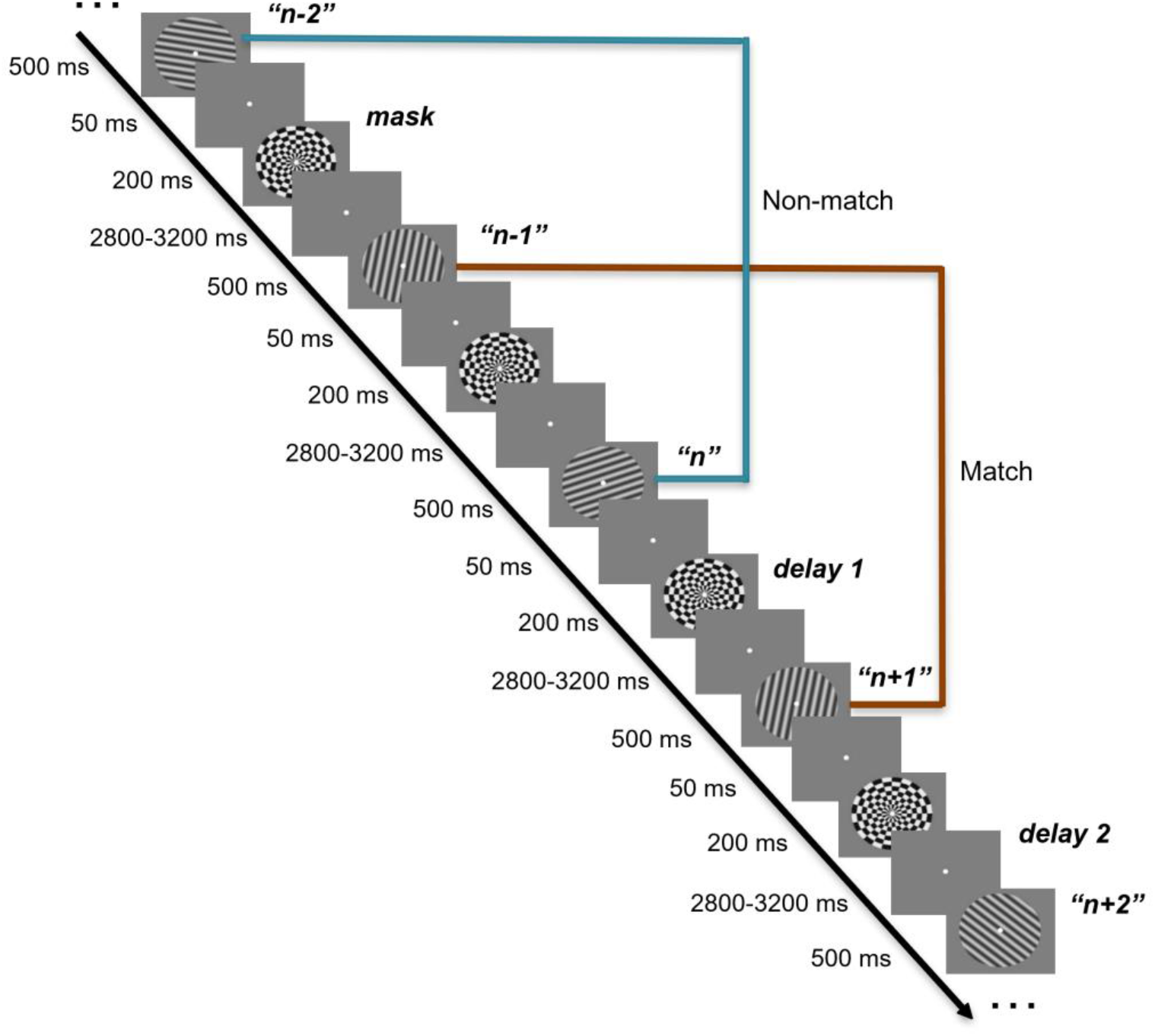
2-back task structure in the Wan et al. (2020) EEG study. The presentation of each stimulus is followed by a 50 ms blank screen, a 200 ms radial checkerboard mask, a variable delay from 2.8 to 3.2 s (only the first 2.8 s, which is common to all stimulus events, was used for analysis), and then the next stimulus was presented, upon which the match vs. non-match response is to be made.

**Figure 2.**
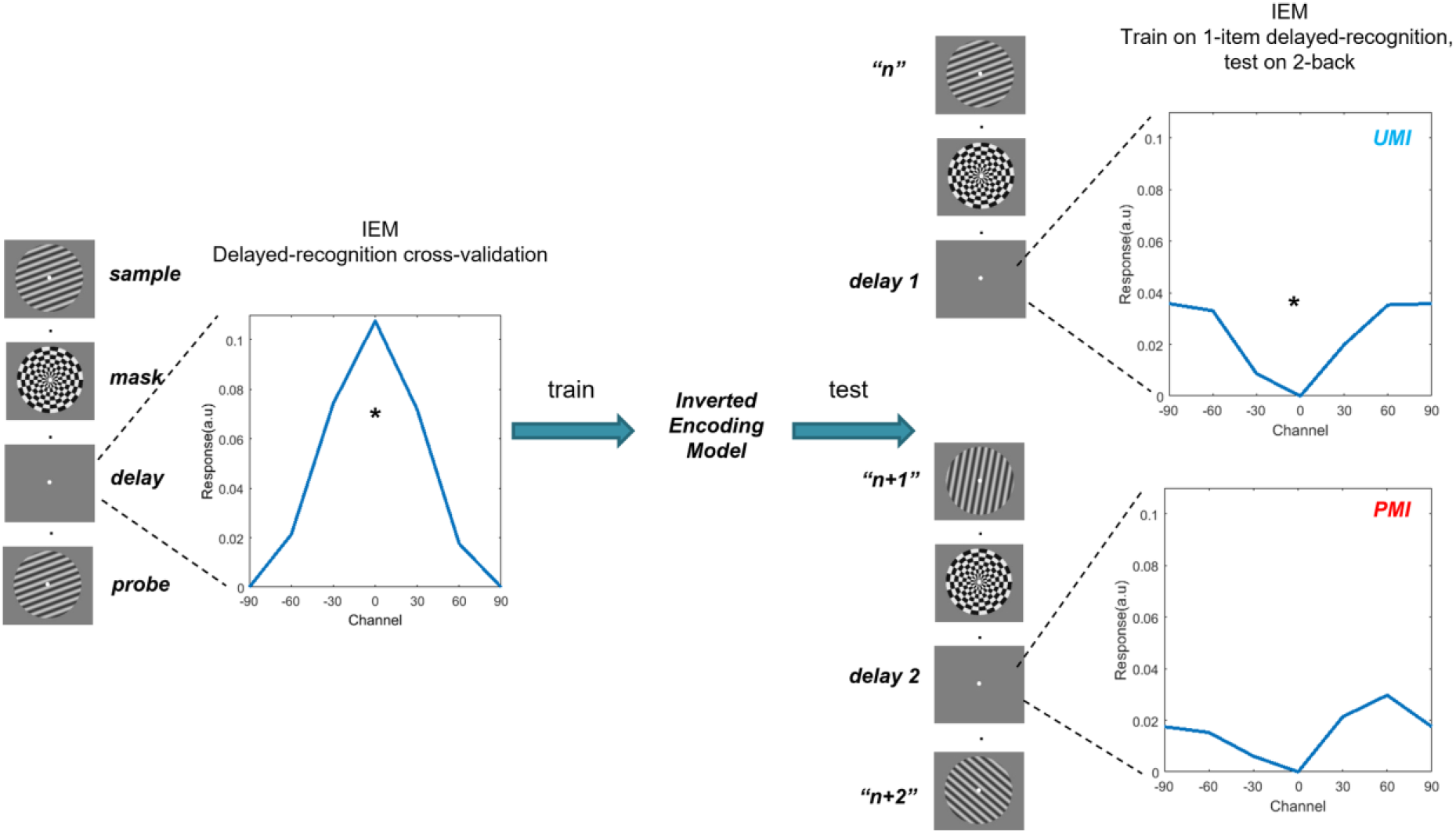
IEM reconstruction of the 2-back task. In IEM, voltage from each EEG electrode is construed as a weighted sum of responses from six orientation channels (modelled by a half-wave-rectified sinusoid raised to the 6^th^ power), each tuned to a specific stimulus orientation, comprising the basis set. Left panel: IEM reconstruction of the stimulus during the delay in a separate one-item delayed-recognition task. This model was used to reconstruct the stimulus in the 2-back task. Right panel: Concatenation of the item *n* and item *n + 1* stimulus events to form a trial, across which *n* transitions from probe to UMI to PMI in the 2-back. On the right are IEM reconstructions corresponding to the two 2 s windows centered in the 2.8 s post-mask ISIs before and after item *n + 1*, respectively. “*” indicates *p* < .01 (two-tailed *t* test), FDR-corrected for multiple comparisons. As the figure shows, IEM reconstruction of stimulus *n* is “flipped” relative to the training data (IEM reconstruction from delayed recognition) when it is a UMI and transformed again when its status becomes PMI, demonstrating priority-based remapping. For delayed-recognition IEM reconstruction (940 – 1040 ms from stimulus onset), *t*(41) = 4.12, *p* < 0.001. For UMI reconstruction of 2-back (−2400 – −400 ms relative to *n + 1* onset), *t*(41) = −3.02, *p* = 0.009; for PMI reconstruction of 2-back (1150 – 3150 ms from *n + 1* onset), *t*(41) = −1.60, *p* = 0.117.

**Figure 3.**
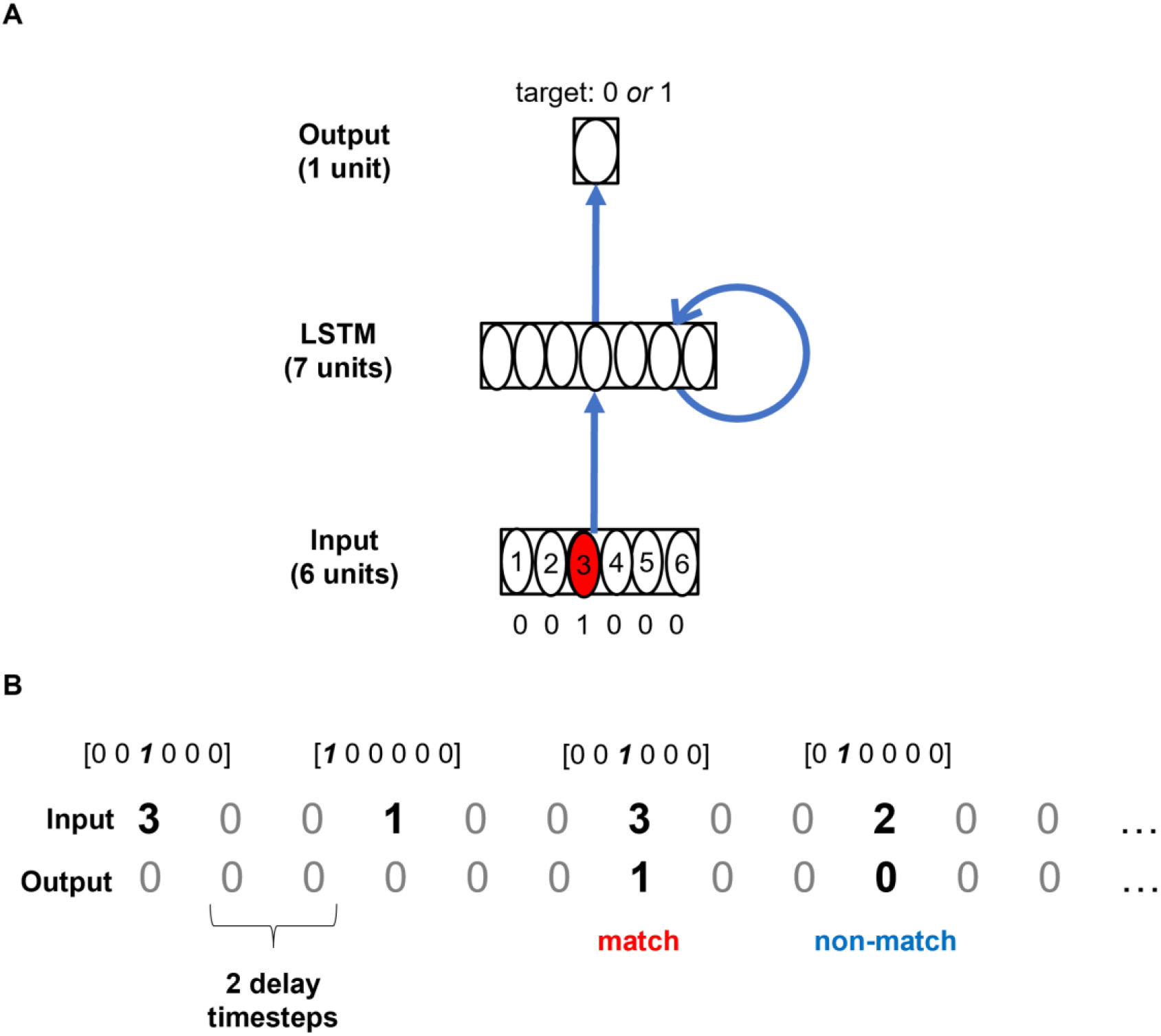
RNN model architecture. (A) One-hot vectors corresponding to each of the 6 stimulus types are fed into the input layer, which projects to an LSTM layer with 7 hidden units. This hidden layer in turn projects to an output unit with a binary target activation (0 = non-match, 1= match). (B) Example input and target output sequences. Two delay timesteps were installed after each stimulus presentation timestep to emulate the delay period in the 2-back EEG task.

Two computational models offer some insight into priority-based remapping. One model, by Lorenc and colleagues (2020), was designed to account for a similar flipped IEM reconstruction observed in an fMRI study using a retrocuing task. This approach was inspired by evidence from monkeys performing WM tasks, in which top-down signals from FEF were shown to alter several receptive field properties of neurons in extrastriate visual areas V4 and MT (Merrikhi et al., 2017). They created simulated data for training IEMs using the basis set that was employed for IEM reconstructions on empirical data, and subsequently created a test dataset where the basis function parameters for memory strength, gain, receptive field width, and receptive field centers were varied. These parameters were then fitted to experimental data. Although this model reasonably reproduced the flipping of IEM reconstructions, its mechanistic interpretation was equivocal because multiple solutions fit the data similarly well (i.e., width modulation versus memory strength + gain modulation). Additionally, its implications for the PMI-to-UMI transition are unclear, because the simulated “flipped” IEM reconstruction was obtained from the time period when the stimulus was no longer required for the task. A second model, from Manohar and colleagues (2019), simulated working memory performance in a network comprised of hard-coded feature-selective units and a pool of freely conjunctive units that can form a plastic attractor to keep one item, a PMI, in a state of elevated activity. When attention shifted away from an item (making it a UMI), it remained briefly encoded in a residual pattern of strengthened connections, and, under some conditions, inhibition from activity in other parts of the network produced an “inverted” representation of UMI. Although this model successfully reproduced other empirical findings using simulated data, such as the temporary reactivation of the UMI by a nonspecific pulse of excitation, it was not used to account for empirical neural data. From the perspective of the framework of Marr and Poggio (1976), both of the models reviewed above were intended to address the phenomenon of opposite results between UMI and PMI at the implementational level (e.g., *why does the IEM reconstruction flip?*). Our interest in this report, however, is not to understand how different conditions might influence the behavior of MVPA or IEM. Rather, our interest is at the algorithmic level of analysis: Are shifts in priority status accompanied by systematic transformations of neural representation? Thus, although MVPA and IEM produced the results that gave rise to the priority-based remapping hypothesis, are poorly suited to evaluating it, because they don’t permit direct measures of neural representation (Gardner & Liu, 2019; Liu et al., 2018; Sprague et al., 2018, 2019).

To address this question, we turned to artificial neural networks (ANNs), which have been playing an increasingly prominent role in providing mechanistic insights into, and generating novel hypotheses of, phenomena in cognition and neuroscience (Kell & McDermott, 2019; Mante et al., 2013; Richards et al., 2019; Sussillo et al., 2015; Yang et al., 2019). In the current work, we use recurrent neural networks (RNNs) with an LSTM architecture (Hochreiter & Schmidhuber, 1997) to perform a 2-back WM task modeled on Wan et al., (2020). LSTMs can generate flexible behavior guided by long range temporal dependencies, and can solve complex tasks such as speech recognition (Graves et al., 2013) and machine translation (Sutskever et al., 2014). Moreover, LSTM might be a good model for WM tasks due to its gating-based architecture, reminiscent of the cortico-striatal mechanisms believed to gate information into and out of WM (Chatham & Badre, 2015; O’Reilly & Frank, 2006).

Our approach was to train RNNs to perform the 2-back task, then first use Principal Component Analysis (PCA) of the activity of the RNN’s hidden layer to visualize its representational dynamics. This revealed a smooth rotational transformation of stimulus representations over the course of the trial (Figure 4). This trajectory was consistent with what would be expected of a series of priority-based transformations of representational formats as stimuli transitioned functional roles from memory probe to UMI to PMI. However, PCA does not allow for the isolation and quantification of variation attributable to specific task dimensions (of particular interest here, priority status and the match/nonmatch decision). Therefore, we treated these observations as a hypothesis-generating step, and carried out two additional sets of analyses. First, we established the validity of these hypotheses in the RNN data by submitting the data to demixed Principal Component Analysis (dPCA; Kobak et al., 2016) -- a procedure that allowed for the identification of dimensionality-reduced subspaces specific to the probe, UMI, and PMI states of representation, as well as one specific to the decision – then quantifying the geometric relations of these subspaces to one another as well as the temporal dynamics of the representational geometry within these subspaces. This yielded the final a priori, quantitative hypotheses that we tested with the EEG data from Wan et al. (2020). The results of these hypothesis tests provided novel insights about priority-based transformations of stimulus information that are carried out by the human brain.

**Figure 4.**
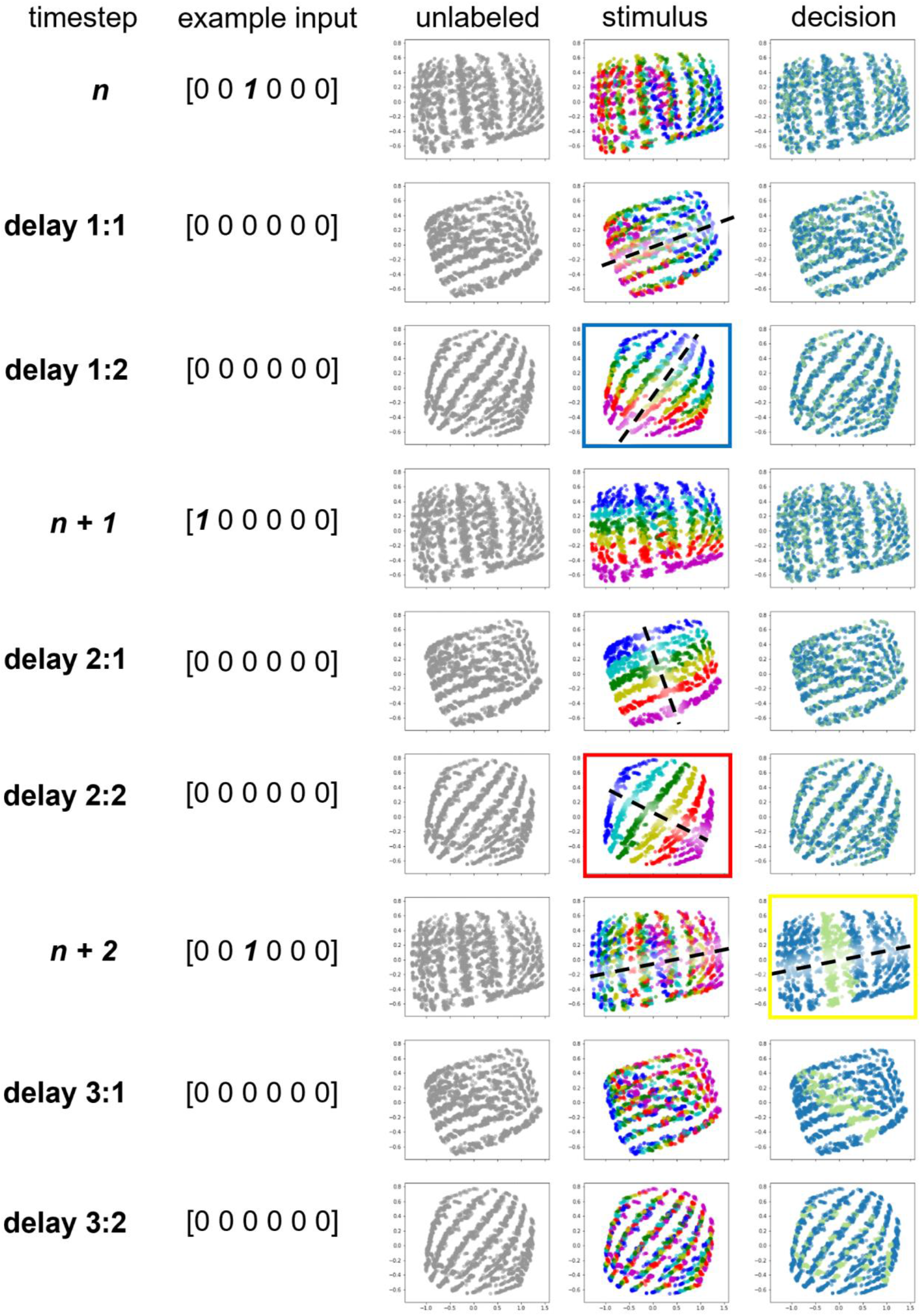
PCA visualization of LSTM hidden layer activity of an example network (#7). Shown is a 9-timestep time course of the 2-back task, running from stimulus *n* to *delay 3:2*. Column 1 and 2: timestep labels and example input vectors. Column 3: Each dot in the figure indicates the representation of stimulus *n*. Column 4: Same as Column 3 but now each color corresponds to one of the six stimulus types, and the black dashed line illustrates the “schematic” stimulus coding direction. Column 6: Same as Column 5 except that the colors now correspond to an item’s status for the *n*-to-*n + 2* comparison that occurs at timestep *n + 2* (green: match trials, blue: non-match trials). Black dashed line at timestep *n + 2* illustrates the decision-based structure. As can be seen in Column 5, the stimulus coding direction rotates counterclockwise (in the image plane) over time such that it becomes “perpendicular” to the decision structure at timestep *n + 1* and aligns with it at timestep *n + 2*.

## Methods

### Behavioral task

In each experimental block of the 2-back working memory task, both human subjects (*N* = 42) and RNNs (*N* = 10) were serially presented a sequence of stimuli drawn from a closed set of six different identities (128-stimulus blocks for humans, 20-stimulus blocks for RNNs). The task was to indicate, for each stimulus, whether or not it matched the identity of the stimulus that had been presented 2 positions earlier in the series. Each EEG subject performed 4 blocks and each RNN performed 200 blocks.

### Recurrent neural network (RNN) model

#### RNN architecture

Ten RNNs with an LSTM architecture were trained and simulated using the Python-based machine learning package PyTorch. Initially, networks consisted of 6 “orientation”-selective input neurons and 7 LSTM hidden units (as provided by default in PyTorch), which were linearly rectified and linearly read out to a single output neuron (Figure 3). We employed the linear rectification to emulate the nonlinear relationship between subjects’ memory representations (as measured by EEG) and their decision outputs. Networks with other numbers of hidden units (up to 256) gave qualitatively similar results. Initially we chose to use 7 units because they are few enough to solve the task, and the network solutions (as evaluated by representational dynamics from the PCA visualization) were the most consistent across training instances). Subsequently, we repeated the procedure with RNNs with 60 LSTM hidden units, to match the dimensionality of our EEG data.

#### Stimuli

The identity of each stimulus presented to the network was denoted by an integer randomly generated between 1 and 6. The stimulus input took the form of a one-hot vector, with only the unit corresponding to the stimulus identity activated (e.g., [0, 0, 1, 0, 0, 0] for stimulus #3; we also explored RNNs trained on metrically varying input vectors following the basis function used to build IEMs in Wan et al. (2020), and these yielded similar results, see Supplementary Materials S1). To simulate the delay period in the human task, we installed 2 “delay” timesteps following the presentation of each stimulus (with an input of [0, 0, 0, 0, 0, 0]; no delay timesteps after the last stimulus in the sequence). A “stimulus event” consisted of the presentation of stimulus *n* and its following two delay timesteps. To evaluate the UMI-to-PMI representational transition of stimulus *n*, we refer to the concatenation of each two consecutive “stimulus events” as a “trial”.

The output unit took on a value at each timestep, with target output values of 0 during each delay timestep, of 0 for stimulus presentation timesteps presenting a non-matching stimulus (equivalent to withholding a response on a target-detection task), and of 1 for stimulus presentation timesteps presenting a matching stimulus (i.e., a stimulus matching the item presented two items previously). Each block comprised 18 trials (because no delay period followed stimulus #20; the last trial contained stimulus #18 and #19), and only 16 trials were analyzed (because the first two stimulus events had no target outputs: not enough stimuli preceded them to have a match/non-match decision). We generated 200 random stimulus sequences for training the RNNs and 200 random sequences for testing the trained networks. Because the human 2-back task had a ratio of 1:2 between match and non-match trials, we generated random sequences that satisfied the criterion that each sequence had to contain at least 5 match trials. The outcome was that training sequences had an average of 5.55 match trials (*SD* = 0.78) and testing sequences an average of 5.46 match trials (*SD* = 0.70).

#### RNN training and testing

Unit activity of the RNNs was initialized with 0, and weights and biases were initialized with random values. The RNNs were trained using the Adam stochastic gradient descent (SGD) algorithm for 5000 iterations (Kingma & Ba, 2017; learning rate = 10^-3^). In each iteration, a batch of 20 sequences was randomly selected (with replacement) from the 200 training sequences. The loss function minimized was the mean squared error between output activity and target output across all timesteps and sequences.

After 5000 iterations of training, RNNs were tested on an independently sampled set of 200 stimulus sequences to assess generalization. The network’s performance accuracy was calculated as the percentage of trials (across all 200 sequences in the test set) on which the network made a correct response, where a response was deemed correct if the absolute difference between the activation of the output neuron and the target output was smaller than 0.5. For 7 hidden-unit networks, we set a criterion level of performance accuracy of 99.5%. Because individual networks revealed very similar network behavior and representational dynamics, we judged that successful training of 10 networks would be sufficient, and total of 12 networks were trained to achieve this subjectively determined number (i.e., 2 networks discarded). All RNNs trained had the same architecture, hyperparameters and training/testing sequences. The only thing that differs across networks is the random initialization of the RNN weights. For analyses, the activity timeseries of the LSTM hidden layer units from all 3200 trials (16 trials x 200 sequences) in the training data set were extracted for subsequent analyses.

After analysis of the 10 successfully trained 7 hidden-unit networks, we repeated these training procedures and trained 10 RNNs with 60 units in the LSTM layer (batch size = 20, learning rate = 10^-3^, 1500 iterations), so as to generate RNN data matching the dimensionality of our EEG data sets.

#### PCA visualization of the LSTM layer activity

We extracted from each network the activity of the 7 hidden units in the LSTM layer from all 200 training sequences and used Principal Component Analysis (PCA) to project these 7-dimensional activity patterns onto the top two dimensions accounting for the most variance across all training sequences and timesteps. We then visualized each stimulus *n*’s transition from probe to UMI to PMI within this subspace by plotting the dimensionality-reduced activity across the 9-timestep time course of a trial. These 9 timesteps comprised the presentations of stimulus *n, n + 1, n + 2* and the delay timesteps that followed each (i.e., *delay 1:1 and delay 1; delay 2:1 and delay 2:2; and delay 3:1 and delay 3:2*; Figure 4, “unlabeled” column). (Note that, once a decision has been made about item *n + 2*, item *n* is no longer relevant for the task, and so the *delay 3:1* and *delay 3:2* timesteps illustrate the evolution of the representational structure of *n* after it has been “dropped from WM”.) To see how the representation of stimulus *n* evolves as it transitions from being a UMI to a PMI, we colored the activity patterns according to the identity of stimulus *n* (Figure 4, “stimulus” column). As explained in the Introduction, the memory of stimulus *n* is a UMI during the delay period after the presentation of stimulus *n* (i.e., during *delay 1:1 and delay 1:2*; because it is not needed for the upcoming *n – 1*-to-*n + 1* comparison), then becomes a PMI during the delay period after the presentation of stimulus *n + 1* (i.e., during *delay 2:1 and delay 2:2*; in preparation for the imminent comparison with *n + 2*). We focused on the *delay 1:2* and *delay 2:2* timesteps (highlighted by blue and red squares) to characterize the UMI-to-PMI representational transformation. To visualize the representation of decision, we re-plotted the same activity patterns but colored them according to the correct response to the *n*-to-*n + 2* comparison when n + 2 was presented (Figure 4, “decision” column).

### WM-specific dimensionality reduction via dPCA

Demixed principal component analysis (Kobak et al., 2016) was employed to identify dimensions of RNN and EEG activity relevant to the stimulus representation in WM. Unlike PCA, which seeks dimensions that maximize the total variance of the data regardless of task variables, dPCA allows one to parse out dimensions of variability specific to certain task variables (e.g., stimulus identity, decision). Given a task variable of interest, the dPCA algorithm does this by grouping neural activity patterns according to this variable and identifying dimensions that maximize between-group variance and minimize within-group variance. Here, we used this method to identify dimensions of activity that were strongly modulated by the identity of the UMI or PMI during the delay period.

To visualize the UMI neural representation, we first sought to identify dimensions along which neural activity was relatively stable over the delay period and also strongly dependent on the UMI stimulus identity. We thus estimated the UMI demixed Principal Components (dPC’s) by minimize the following loss function:

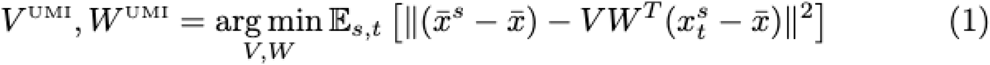

where 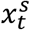 is the neural activity at time *t* within the delay period averaged over all trials in which stimulus *s* (*s* in [1, 2, 3, 4, 5, 6]) was the UMI (trial averaging was necessary to average away noise), 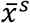 is its temporal mean over the delay period, and 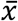 is the global mean over all trials and delay period timepoints. We used *delay 1:2* (for UMI) and *delay 2:2* (for PMI) for RNN and the second half of the delays (−1400ms to 0ms (for UMI) 2150ms to 3550ms (for PMI) relative to stimulus N+1 onset) to find stimulus-specific dimensions. The N x D matrix *V^UMI^* is the so-called “encoding” weight matrix and the N x D matrix *W^UMI^* is the “decoding” weight matrix, used for dimensionality reduction, where N is the dimensionality of the neural activity (7 for RNN, 60 for RNN and 60 for EEG data). This optimization problem is called a reduced-rank regression problem, and admits a closed-form solution (Kobak et al., 2016). Because there are only 6 different UMI stimuli (and thus 6 different 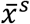 vectors), only up to D=5 dPC’s can be computed in this way (since the ordinary least-squares solution has rank 5). As in PCA, these dPC’s can be ordered in terms of the amount of variance they explain in the data.

Because the task variable used for dPCA was the UMI stimulus, we call these dPC’s the UMI dPC’s, and call the subspace spanned by corresponding encoding vectors, *V^UMI^*, the UMI subspace. We also extracted PMI dPC’s, *V^UMI^*, *W^UMI^*, and a PMI subspace by exactly repeating the above operation but with the index *s* now indexing the PMI stimulus identity.

For visualization purposes we used D=2 dPC’s to obtain 2-dimensional projections, 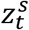 of the neural activity. These projections were computed using the dPC decoders,

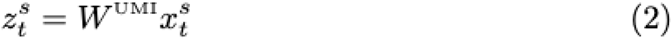

It is these 2-dimensional vectors that are plotted in Figure 5 for the simulated RNN data (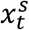 is the internal LSTM state vector) and the EEG data (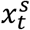 is the vector of EEG signals recorded at each channel; Figure 6).

**Figure 5.**
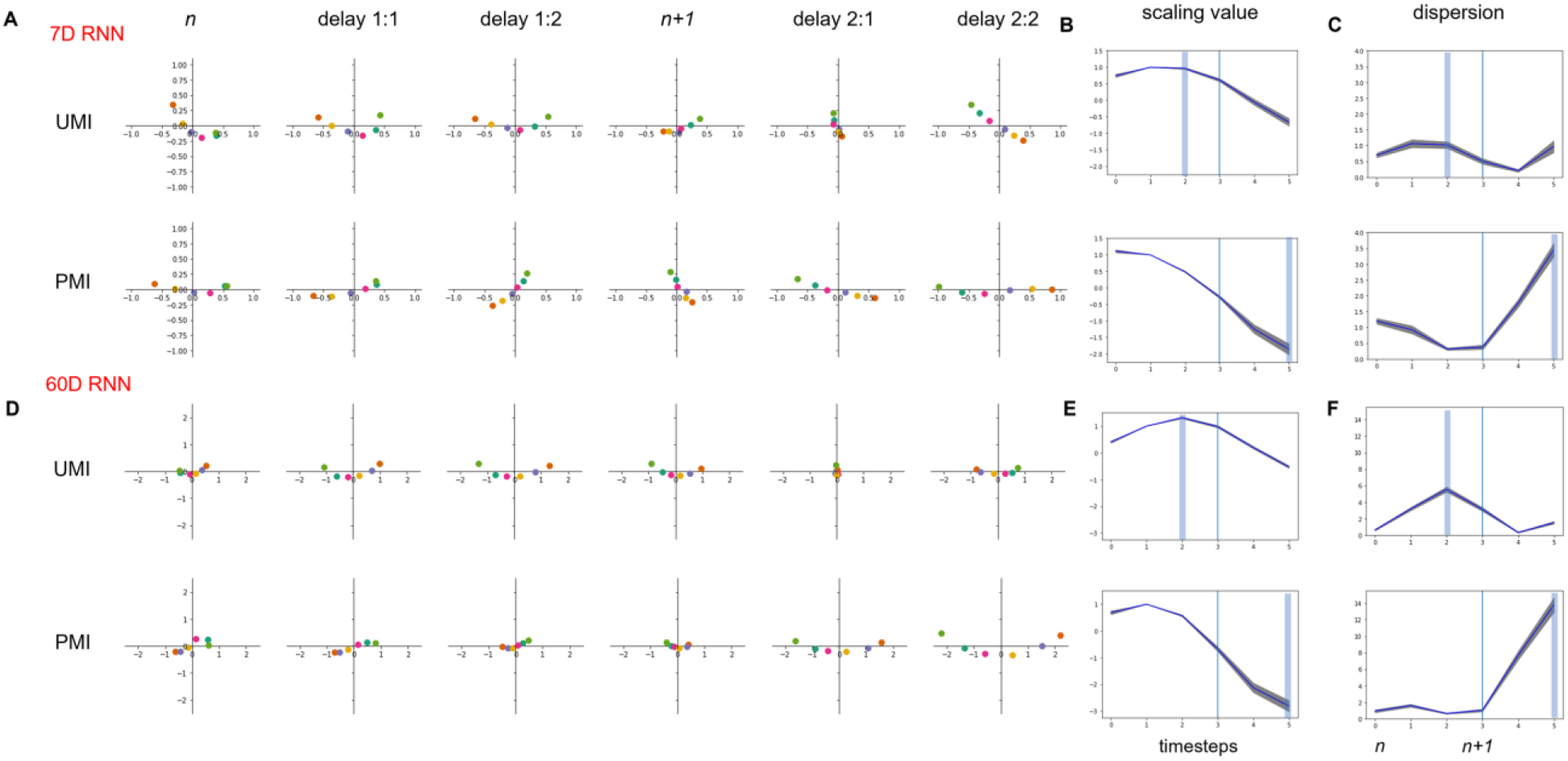
(A) the stimulus averages of RNN hidden layer activity projected into UMI and PMI subspaces over the course of a trial (stimulus *n* to *delay 2:2*) for an example 7-unit network. On the right, the time courses of (B) scaling value and (C) dispersion metric that capture the representational transformation. Blue vertical line indicates the presentation of stimulus *n + 1*. Light blue shading shows the timesteps that were used to identify the dPCs. The gray shading around the curve shows standard error from all 10 trained RNNs. (D, E, F) Same as (A, B, C) but for the 60-unit RNNs.

**Figure 6.**
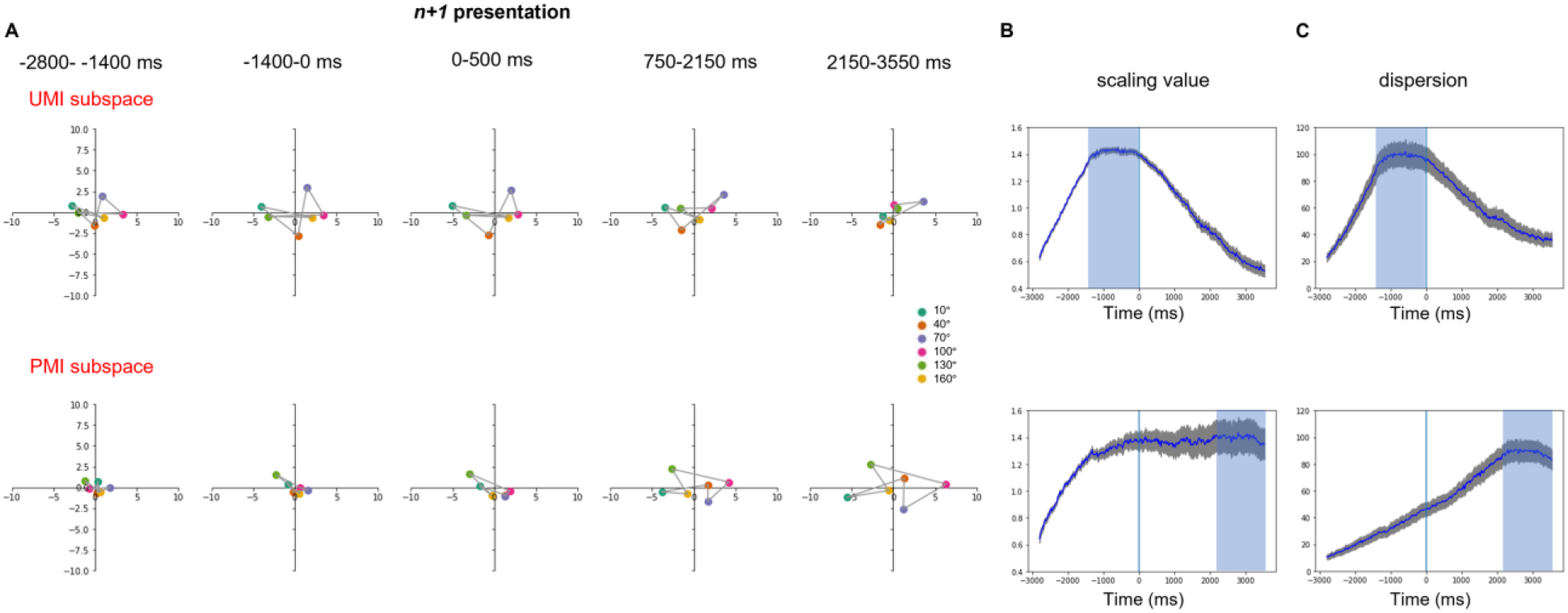
(A) the stimulus averages of EEG signal projected into UMI and PMI subspaces over a 2-delay time course (−2800ms to 3550ms relative to stimulus *n + 1* onset) for an example subject. Data points of adjacent stimulus angles are connected by a gray line. On the right, the time courses of (B) scaling value and (C) dispersion metric that capture the representational transformation. Blue vertical line indicates the onset of stimulus *n + 1*. Light blue shading shows the time windows that were used to identify the dPCs. The gray shading around the curve shows standard error from all 42 EEG subjects.

For estimating stimulus and decision subspaces (Figure 7), a slightly different loss function was employed. We sought to capture dimensions of decision variability across different stimuli. We expected that decision variability might depend on the stimulus identity of the probe, so in our dPCA analysis we aimed to capture decision related fluctuations at each stimulus and timepoint. For stimulus dPC’s, we thus minimized the following loss:

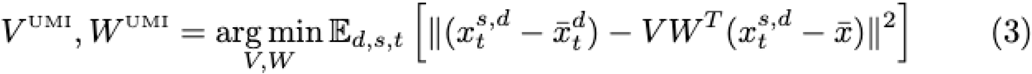

where 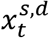 is the is the neural activity at time *t* within the second half of the delay period averaged over all trials in which stimulus *s* was the UMI and response *d* (“match” or “non-match”) was the decision made by the subject/RNN, 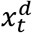 is its mean over stimuli, and 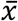 is again the global mean over all trials and delay period timepoints. PMI dPC’s were computed by replacing the *s* index with an index of the PMI stimulus. Decision dPC’s were similarly computed by minimizing the following loss:

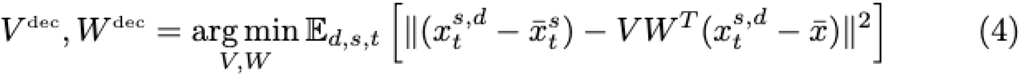

**Figure 7.**
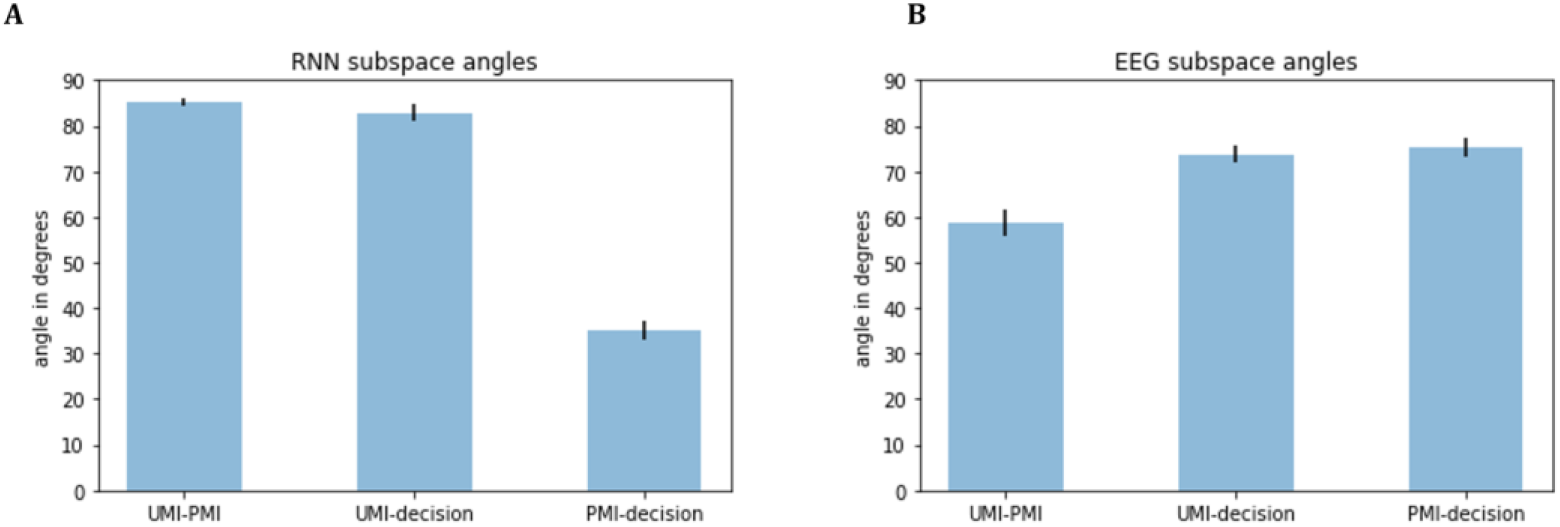
Angles between UMI, PMI and decision subspaces for (A) RNN and (B) EEG data. Black bars show standard error.

Here, 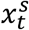 is the neural activity averaged over all decisions. We again used D=2, in accordance with previous analyses of WM subspaces (Panichello & Buschman, 2021). See “UMI/PMI/decision subspace analysis” section below on how the relationships between these subspaces were then quantified.

Percent variance explained calculations were performed as follows. Percent global variance explained by the *i*th dPC, *w_i_* (i.e. the *i*th row of the decoder matrix *W*), was calculated using the corresponding column *v_i_* from the encoder matrix *V* by

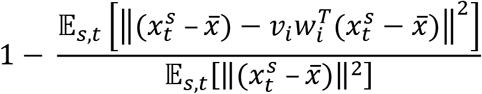

The percent stimulus variance explained was defined as

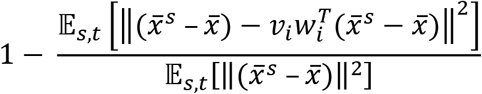

### Characterizing the dynamics of the UMI-to-PMI transformation

To characterize the continuous dynamics of the UMI-to-PMI rotational transformation in stimulus-relevant dimensions, we quantified the evolving geometry of the UMI and PMI representations visualized in Figure 5 and 6 with two different scalar metrics.

The first metric quantified how dispersed these points were from each other:

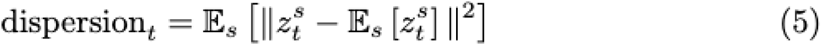

In Figure 5C and 6C we plot this dispersion metric as a function of time, for both the low-dimensional projections, 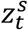 computed from the UMI dPC’s and PMI dPC’s calculated according to equations 1 and 2.

The second metric quantified the change in the UMI/PMI representation over the trial relative to time-averaged representation during the first half of the first delay. This change was quantified using a scalar,

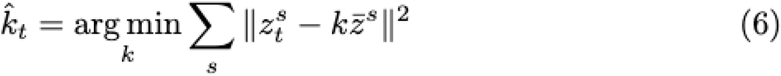

where 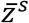 here refers to the time-averaged projection over the timesteps during the first half of the first delay (*delay 1:1* for RNN and −2800 to −1400 ms relative to stimulus *n + 1* onset for EEG). In Figure 5B and 6B we plot these best-fitting scalars as a function of time, using both the low-dimensional projections, 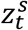, computed from the UMI dPC’s and PMI dPC’s calculated according to equations 1 and 2.

### UMI/PMI/decision subspace analysis

To quantify the relationship between UMI, PMI and decision subspaces calculated from equations 3 and 4, we used a metric developed by Panichello and Buschman (2021). This metric measures the alignment between corresponding pairs of dPC encoding vectors as follows:

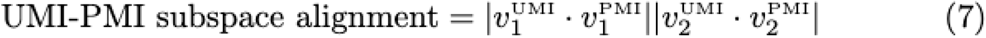

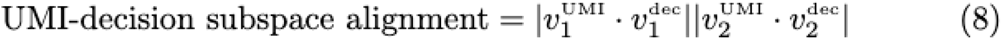

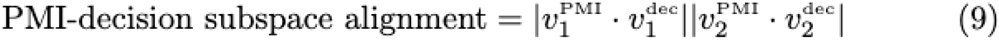

where the dot denotes the Euclidean dot product, and the bars denote absolute value. Here, 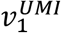, 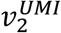 are the 1^st^ and 2^nd^ UMI dPC encoding vectors, i.e. the two columns of the N x 2 matrix *V^UMI^*. The analogous definition holds for the PMI and decision dPC’s: 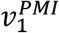, 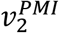 are the columns of *V^PMI^*; 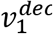, 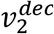 are the two columns of *V^dec^*. Note that under the standard dPCA formulation used by Kobak et al. (2016) and used here, the encoding vectors are all norm 1. These dot products can therefore be interpreted as cosines of angles between the pairs of vectors, and the subspace alignment metric can be interpreted as a product of two cosines.

To turn this metric into an angle, we took the inverse cosine of each alignment metric in equations 7, 8, 9. These are the angles plotted in Figure 7.

### EEG dataset

The experimental protocol for the Wan et al. (2020) EEG study (the data from which was analyzed in this paper), along with the informed consent form, was approved by the University of Wisconsin–Madison Health Institutional Review Board (protocol no. 2016-0500). Prior to each experimental session, informed consent was obtained by lab personnel listed on the IRB-approved protocol.

60-channel EEG data were acquired and preprocessed as per procedures described in Wan et al. (2020). Raw EEG voltages were used for all analyses. Because data from the pilot and replication experiments from Wan et al. (2020) yielded very similar IEM reconstruction results, they were combined to yield a dataset of 42 subjects. As is the case with the RNN data, after excluding the first two stimuli from each block there were 126 stimulus events and hence 125 trials per block (Figure 2). Each stimulus event (stimulus presentation followed by a delay) lasted 3550 ms. A third of the trials in each block were ‘match’ trials and the other two thirds were ‘non-match’ trials. EEG data from all trials (both correct and incorrect) were included in the analyses. For each stimulus *n*, during the delay period after its onset, stimulus *n – 1* had the status of PMI and *n* had the status of UMI.

## Results

### Behavioral results of EEG study

Mean accuracy was 86.1% (*SD* = 5.6%), mean *d’* was 2.40 (*SD* = 0.65), and mean response time was 0.82 s (*SD* = 0.18 s).

### PCA of LSTM activity patterns

PCA was carried out on the 7D LSTM hidden layer activity from the training data, and the resultant dimension-reduced activity from all 3200 trials projected onto the 2D-space constructed by the first 2 principal components (Figure 4, “unlabeled” column). This revealed that representations tended to cluster into band-like manifolds that appeared to rotate over the course of the trial (i.e., from timestep *n* to timestep *n + 2*). Next, to get a sense of the stimulus representational structure and how it evolves over time, we colored the data points for each trial according to the identity of stimulus *n* (Figure 4, “stimulus” column). This revealed that, across trials, stimulus representations were organized into stimulus-specific “stripes” that at some timesteps cut across the band-like manifolds (*delay 1:1* and *delay 1:2*), and at others were perfectly overlaid on them (*delay 2:1* and *delay 2:2*). These “stripes” thus defined a stimulus-coding axis. (That is, a stimulus’s identity can be read out based on its location along this axis. A schematic illustration of this axis is superimposed on some of the timesteps from Figure 4, “stimulus” column, with a black dotted line.) It is noteworthy that, at timestep *n + 2*, the configuration of individual trials is different than at timestep *n*. This reflects that fact that items serve different functions at these two timesteps – *probe* at timestep *n* and *memorandum* at timestep *n + 2*. Indeed, if one were to re-color timestep *n + 2* according to stimulus *n + 2*’s identity, this frame would be identical to the configuration of stimulus *n* at timestep *n*, which means that *n + 2* and *n* are in opposite locations in PCA space (e.g., in Figure 4, the azure-colored stimulus trials occupying the right side of PCA space at timestep *n* are on the left side of the space at timestep *n + 2*).

Finally, to get a sense of the decision representational structure, we colored each data point according to whether or not it would require a response of “match” at the end of the trial (i.e., at timestep *n + 2*; Figure 4, “decision” column). This revealed that, when *n* is compared with *n + 2*, trials requiring a “match” response converged onto the two central band-like manifolds, whereas non-match trials the flanking manifolds. This organization thus defined a decision-coding axis, in that the output required for a stimulus can be read out based on its location along this axis. A schematic illustration of this axis is superimposed on some of the timesteps from Figure 4, “stimulus” column, with a black dotted line).

Over the course of a trial, *n*’s stimulus-specific axis appeared to rotate counterclockwise (in the PCA plane) as it transitioned from UMI (during *delay 1:1 and delay 1:2*) to PMI (during *delay 2:1 and delay 2:2*). This likely reflects, in part, transitions between the functional roles of probe (timestep *n*), then UMI, then PMI. Thus, we can hypothesize the following functional account of the representational trajectory through a trial of, say, an azure-colored stimulus from Figure 4. At timestep *n*, its representational structure puts it on one of the central bands if it matches item *n – 2* (and therefore elicits an output of [1]), or on a band to the right of center if it does not match item *n – 2*. These two locations align with the decision-coding axis. Next, as it acquires the functional status of UMI, it transitions to a configuration that is not compatible with decision-making, as evidenced by the fact that every azure stimulus is located along a “stripe” that is parallel to the decision-coding axis (stated another way, the stimulus-coding axis at timestep *n + 1* is orthogonal to the decision-coding axis). During *delay 2:1* and *delay 2:2* the item’s representation continues to rotate in the same counterclockwise direction, trajectory that brings it back into alignment with the decision axis, but now on the “opposite side” of the PCA space, reflecting the fact that it is a PMI. (I.e., for azure items, probes cluster on the right side of PCA space, PMIs on the left side.) At timestep *n + 2*, the band occupied by this item will depend on its match/nonmatch status. From this we can further hypothesize that the function of this rotational trajectory might be to prevent the remembered representation of *n* from influencing the *n – 1* versus *n + 1* decision (at timestep *n + 1*).

Whatever the intuitive appeal of these hypotheses, they can’t be assessed quantitatively because the PCA didn’t allow for the direct comparison of representational geometries between functional states. This is because the PCA had no information about stimulus identity (or any other task variables of potential interest).

### dPCA of LSTM activity patterns

Unlike PCA, dPCA would allow for the identification of subspaces in the data that are specific to stimulus representation. We performed dPCA at multiple timesteps to obtain snapshots of stimulus representational geometry at different points in time. More specifically, it would identify the dimensionality-reduced subspace occupied at timesteps of interest. This would allow us to test quantitatively the hypothesis that, for a given item *n*, its representational format while a UMI is incompatible for readout for the decision about *n + 1*, and that it then transitions into a format that is amenable for readout for the decision about *n + 2*.

#### RNN with 7 LSTM units

To isolate the transformations observed in the PCA (Figure 4), we applied dPCA to the 7D data from the RNNs to identify the top two UMI-selective dPCs (at the *delay 1:2* timestep) and the top two PMI-selective dPCs (at the *delay 2:2* timestep. The first and second dPCs of the UMI subspace accounted for 92.0% and 4.2% of the total stimulus variance of the trial-averaged data, respectively. The first and second dPCs of the PMI subspace accounted for 97.4% and 2.4% of the total stimulus variance, respectively (see Supplementary Materials S2 for additional information). Comparison of the UMI mean (i.e., timestep *delay 1:2* in UMI row of Figure 5A) and the PMI mean (i.e., timestep *delay 2:2* in PMI row of Figure 5A) in the same subspace showed that although both represent stimulus identity along their 1^st^ dPC, they do so in reversed order (i.e., for the network illustrated in Figure 4A, in both subspaces, the ordering along the 1^st^ dPC of *delay 1:2* is *orange-yellow-purple-pink-teal-green*, and the ordering along the 1^st^ dPC of *delay 2:2* is *green-teal-pink-purple-yellow-orange*). (Note that for RNN simulations, stimuli were not metrically related, and so, e.g., the ordering of colors was different for each network’s UMI mean. Nonetheless, for every network this ordering was reversed for PMI mean relative to UMI mean.) Iteratively projecting trial-averaged activity from each timestep on onto these two dPC subspaces suggested that the evolution of stimulus representational format across the trial is such that its projection onto the 1^st^ dPC of the PMI – the axis that is critical for readout of the memory item against which the impending probe is to be compared -- is minimal at timestep *n +1*.

Quantitative elements of the representational trajectory across the trial are captured by the scaling parameter of each timestep relative to a subspace, which captures the rate and direction of change, and the dispersion of means at each timestep, which indexes relative discriminability. Relative to the UMI subspace (i.e., the dPCA of timestep *delay 1:2*), an item’s representational format was relatively stable (i.e., unchanging) for the first half of the trial, then, after timestep *n + 1*, shifted to a steady rate of transformation for the remainder of the trial, with the 0-crossing of the scaling parameter (indicating the reversal of the stimulus coding axis) occurring at timestep *delay 2:1*. Across this trajectory, individual stimulus identities first expanded, then contracted to a minimum dispersion value (i.e., point of least discriminability) at timestep *delay 2:1*, then expanded again (Figure 5C). Relative to the PMI subspace, in contrast, the process of representational transformation began early in the trial -- at timestep *delay 1:1* -- and continued at roughly the same rate for the remainder of the trial. Across this trajectory, stimulus identities first contracted -- achieving their lowest dispersion values at timesteps *delay 1:2* and *n + 1* -- then expanded dramatically across timesteps *delay 2:1* and *delay 2:2*; Figure 5F). Together, these results confirm that an item’s representational transformation across the trial proceeds at a relatively steady rate (consistent with the smooth rotation observed with the PCA (Figure 3)), via a trajectory that minimizes discriminability at timestep *n + 1*, thereby achieving the goal of minimizing the likelihood that the UMI can interfere with the decision about item *n + 1*.

#### RNN with 60 LSTM units

Although the results from the 7D RNN data produced quantitative predictions about the priority-based transformation of information held in WM, their direct applicability to the EEG data from Wan et al. (2020) would be complicated by the difference in dimensionality between the two datasets. Therefore, our next step was to repeat the procedure described up to this point, but with 10 RNNs with 60 LSTM units each. Results with the resultant 60D data would constitute the hypotheses that we would then test with the EEG data from Wan et al. (2020).

10 networks were trained in order to generate 10 that performed the 2-back task at > 99.5% correct. dPCA of the resultant 60D data from the RNNs identified the top two UMI-selective dPCs (at the *delay 1:2* timestep) and the top two PMI-selective dPCs (at the *delay 2:2* timestep). The first and second dPCs of the UMI subspace accounted for 78.7% and 15.1% of the total stimulus variance of the trial-averaged data, respectively. The first and second dPCs of the PMI subspace accounted for 86.1% and 9.9% of the total stimulus variance, respectively (see Supplementary Materials S2 for additional information).

Results with the 60D RNN data also served as the hypotheses that we would test on the EEG data from Wan et al. (2020), and so are organized here in terms of hypothesis.

- *Comparison of UMI mean vs. PMI mean* (i.e., timestep *delay 1:2* of Figure 5D vs. timestep *delay 2:2* of Figure 5D in both UMI and PMI rows). As with the 7D RNN data, ordering of stimulus identity along their first dPCs was reversed.
- *Scaling across the trial, relative to UMI subspace*. The scaling parameter increased at a relatively slow rate from timestep *n* to *delay 1:2*, then decreased for the remainder of the trial, reversing sign timestep *delay 2:1* (Figure 5E).
- *Dispersion across the trial, relative to UMI subspace*. The 60D RNN data showed a pattern of expansion from timestep *n* to timestep *delay 1:2*, then contraction until reaching it minimum dispersion value at timestep *delay 2:1*, before rebounding slightly at timestep *delay 2:2* (Figure 5F).
- *Scaling across the trial, relative to PMI subspace*. The scaling parameter was relatively flat from timestep *n* to *delay 1:2*, then decreased at a steady rate for the remainder of the trial, reversing sign at timestep *n + 1* (Figure 5E).
- *Dispersion across the trial, relative to PMI subspace*. Dispersion value of the 60D representation was low, and unchanging, from timestep *n* to timestep *n + 1*, before rapidly expanding across timesteps *delay 2:1* and *delay 2:2*.
- *Relative orientation of subspaces*. In addition to minimizing discriminability, a second way to minimize the ability of the UMI to interfere with the processing of *n + 1* would be to orient its subspace orthogonal to the probe subspace. Indeed, an angle of 82.87° (*SD* = 5.32°) separated the UMI and decision subspaces. In contrast, the PMI and decision subspaces were separated by a much smaller angle of 35.10° (*SD* = 6.08°), consistent with the prediction from the PCA (Figure 3) that, as the time of presentation of item *n + 2* drew near, the PMI was rotating into an orientation that would facilitate the *n* vs. *n + 2* comparison. The angle separating the UMI and PMI subspaces was 85.09° (*SD* = 2.74°).

### dPCA of EEG activity patterns

The EEG data from Wan et al. (2020) were markedly noisier than the RNN data: The first and second dPCs of the UMI subspace accounted for 45.4% and 23.7% of the total stimulus variance of the trial-averaged data; and the first and second dPCs of the PMI subspace accounted for 46.2% and 23.2% of the total stimulus variance of the trial-averaged data.

#### Hypothesis tests

- *Comparison of UMI mean vs. PMI mean* (Figure 6A). The circular organization of the six stimuli meant that their distribution relative to the 1^st^ dPCs would not be expected to align along their 1^st^ dPC in a manner similar to the RNN data. Nonetheless, inspection of the data from a representative subject (Figure 6A) shows no suggestion of a reversal akin to what was observed in the RNN data.
- *Scaling across the trial, relative to UMI subspace*. The trajectory of the scaling parameter qualitatively matched that from the 60D RNN, increasing from timestep *n* to *delay 1:2*, holding a constant value across timesteps *delay 1:2* and *n + 1*, then decreased for the remainder of the trial. Unlike the RNN data, however, the scaling parameter never reversed sign (Figure 6B).
- *Dispersion across the trial, relative to UMI subspace*. The time course of dispersion was also qualitatively matched to that from the 60D RNN, expanding early in the trial, plateauing across late *delay 1* and the processing of item *n + 1*, then contracting for the remainder of the trial (Figure 6C).
- *Scaling across the trial, relative to PMI subspace*. The trajectory for the EEG data was a steady increase from timestep *n* to *delay 1:2*, after which the scaling parameter was unchanged for the remainder of the trial (Figure 5E). This indicates that stimulus representations begin transforming toward their configuration in the PMI subspace and fully achieve it by epoch *n + 1* (at which time they have UMI status), then maintain this end-state configuration for the remainder of the trial. This trajectory is opposite of the 60D RNN, for which configuration relative to the PMI was unchanging until after delay *1:2*, then rapidly changing across the second half of the trial. Also different from the RNN, the EEG scaling trajectory did not reverse sign.
- *Dispersion across the trial, relative to PMI subspace*. The stimulus representation expanded at a steady rate across almost the entirety of the trial, before asymptoting during *delay 2*. This is also markedly different from the pattern observed with the 60D RNN, for which the dispersion time course closely followed its scaling time course.
- *Relative orientation of subspaces*. An angle of 73.62° (*SD* = 12.73°) separated the UMI and decision subspaces, and an angle of 75.72° (*SD* = 13.93°) separated the PMI and decision subspaces, a finding inconsistent with the pattern observed with the 60D RNN. The angle separating the UMI and PMI subspaces, 60.70°(*SD* = 19.02°), was markedly smaller that for the RNN data.

## Discussion

Results from previous neuroimaging studies have given rise to the idea that representations in working memory undergo a “priority-based remapping” when they obtain the status of UMI (van Loon et al., 2018; Wan et al., 2020; Yu, Teng & Postle, 2020), but the mechanism underlying this transformation was unknown. Here, using neural network modeling and dimensionality reduction techniques, we have identified a transition through representational space that may reflect a general solution to the computational problem of needing to hold information in an accessible state (i.e., “in WM”) but in a manner that won’t influence ongoing behavior. However, noteworthy differences between RNNs and human EEG suggest important differences in implementational specifics, highlighting important questions for future work.

The 2-back task requires information to evolve through three distinct functional states: a probe requiring comparison with the mnemonic representation of item *n – 2* and an overt match/nonmatch report; unprioritized (a state that should minimize interference with the concurrent *n – 1* vs. *n + 1* comparison and report); and prioritized. PCA of hidden-layer activity of RNNs underwent a smooth rotation through 180° of the 2D space defined by the first two PCs. dPCA of RNNs characterized distinct subspaces corresponding to these states, and the trajectories between them. For the 60D RNN data, the UMI and decision subspaces were separated by mean angle of 82.87°, consistent with an orthogonal orientation that would minimize the influence of the UMI on concurrent processing of the probe. By contrast, the PMI and decision subspaces were separated by an angle of just 35.10°, consistent with close alignment that would facilitate comparison of the two.

The organization of these functional subspaces is reminiscent of recent findings from nonhuman primates performing a retrocuing WM task. Subjects first encoded two stimuli – one above fixation and one below -- into WM, then viewed a cue indicating which one to report. Prior to the cue, PCA indicated that the *above* and *below* items were initially represented in subspaces of neural activity separated by a median angle of 79.1°, but that after the cue, the selected item transitioned into a different subspace, and the selected-*from-above* and the selected-from-*below* subspace were closely aligned -- separated by only 20.1°. They interpreted this as a transition of the selected item from a representational format that emphasized the distinction between the two items to a “template” format that abstracted over location (no longer a relevant parameter) and facilitated read-out (Panichello & Buschman, 2021). In our 2-back task, the UMI-to-PMI transition can be understood as the implicit selection of the UMI that occurs after a response is made to item *n + 1*. An important difference between our 2-back task and the retrocuing task of Panichello and Buschman (2021), however, is that their task lacked a UMI state. Rather, after the retrocue, there was no possibility that the uncued item might be needed. Nonetheless, in the PFC, a representation of the uncued item persisted, and its uncued subspace was orthogonal to the template subspace. Therefore, one important question for future work is whether, and if so how, the transition to UMI differs from the transition to no-longer-needed (i.e., “dropping” an item from WM).

Although the EEG data also showed a progression through priority states, only parts of its transformational trajectory resembled that of the RNN data. In particular, human subjects seemed to have recoded item *n* into its UMI and PMI configuration simultaneously, then prepared for *n + 2* by collapsing the UMI subspace during *delay 2*. Perhaps relatedly, whereas UMI and PMI subspaces in the RNN data were separated by an angle of 85° (approaching orthogonality), in the EEG data there were separated by an angle of only 60°, indicating considerable overlap. (Although we currently do not have an understanding of why the PMI subspace evolves differently in EEG relative to RNN, its overlap with the UMI subspace may help explain why, with fMRI and EEG data, multivariate models trained on the PMI can successfully recover information about the UMI (van Loon et al., 2018; Wan et al., 2020; Yu, Teng & Postle., 2020).

It is important to note that the RNN modeling is not intended to simulate EEG data, nor the human brain, which has vastly different structural and functional architecture from our RNNs. For example, because of the relative simplicity of the RNN architecture, and the absence of many sources of noise that are characteristic of EEG (e.g., physiological noise, uncontrolled mental activity, measurement noise), the variability and SNR of the two signals differ markedly. This limits what can be interpreted from direct comparisons between the two sets of results. Nonetheless, an important role for these RNN stimulations has been to establish the validity and interpretability of our approach with dPCA. This, in turn, allowed us to use dPCA to evaluate neural coding in an EEG data set in which multivariate methods failed to find evidence for an active representation of the PMI (Figure 2). It may be this approach, or one like it, will be helpful with other data sets for which the absence of multivariate evidence has led to speculation about putative activity-silent mechanisms in WM.

It is also important to note that the RNNs we simulated have a simple architecture, with a homogeneous LSTM layer, which is, of course, very different from the brain with its heterogeneous patterns of connectivity between neurons with varied functional and structural properties. The RNN simulations of Masse et al. (2019), employing different cell types and explicitly simulating factors like receptor time constants and presynaptic depletion of neurotransmitter, offer one promising example for developing more biologically plausible models. Also missing from our RNN architecture is an explicit source of control, such as that exerted by prefrontal and posterior parietal circuits in the mammalian brain. Through extensive training, our RNNs gradually learned to adjust their connection weights so as to achieve a high level of performance, but this was only possible because each item presented to the network always followed the same functional trajectory (probe, then UMI, then PMI). A hallmark of WM in the real world is the ability to flexibly respond to unpredictable changes in environmental exigencies. Thus, an important future goal will be to extend the present work to a network with separate modules with different connectivity patterns and governed by different learning rules (e.g., Kruijne et al., 2020; O’Reilly & Frank, 2006), and to a task that requires truly flexible behavior.

Our work complements extant models of attentional prioritization in WM. First, it sheds light on the prioritization mechanisms of a continuous-performance WM task (2-back), a design that has recently received less attention than tasks employing retrocuing. Second, compared with the aforementioned computational accounts (Lorenc et al., 2020; Manohar et al., 2019), our use of dPCA provides a data-driven dimensionality reduction approach that does not make assumptions about the representational structure of stimuli. This allows one to examine the unmodeled structure of stimuli in the representational space. Third, our dPCA analyses were applied on a subject-by-subject basis, without assuming that the same representational and/or computational scheme is employed across individuals. Indeed, recent research has shown that representational biases of stimulus features vary among individuals in higher-order brain areas (Gong & Liu, 2020). Therefore, this approach may be helpful for explaining individual differences across many types of cognition.

To conclude, we used ANN simulations to validate the idea, at the level of representational codes, that shifts of priority status trigger the transformation of stimulus representations in WM. Applying dimensionality reduction to activity patterns from LSTM units in RNNs revealed the organization of functionally specific subspaces, and the trajectories between them. This approach translated to EEG data from subjects performing the same task, revealing similarities and differences between human and machine, and highlighting fruitful directions for future research.

## Data Availability

All processed human EEG data, code, network training sets and trained networks are available at https://osf.io/sgqvn/ (DOI 10.17605/OSF.IO/SGQVN) on Open Science Framework.

## Author Contributions

Conceptualization: Q.W., B.R.P.

Data Curation: Q.W.

Formal Analysis: Q.W., J.A.M.

Funding Acquisition: B.R.P.

Investigation: Q.W., J.A.M., B.R.P.

Methodology: Q.W., J.A.M.

Project Administration: Q.W.

Resources: *Not Applicable*

Software: Q.W., J.A.M.

Supervision: B.R.P.

Validation: Q.W., J.A.M.

Visualization: Q.W., J.A.M.

Writing – Original Draft Preparation: Q.W., J.A.M., B.R.P.

Writing – Review & Editing: Q.W., J.A.M., B.R.P.

## Acknowledgements

We thank Drs. Yuri Saalmann, Joseph Austerweil, Timothy Rogers, Jacqueline Fulvio, Qing Yu and Mohsen Afrasiabi for helpful discussion and critical feedback.

## Supplementary Materials

**Figure S1.**
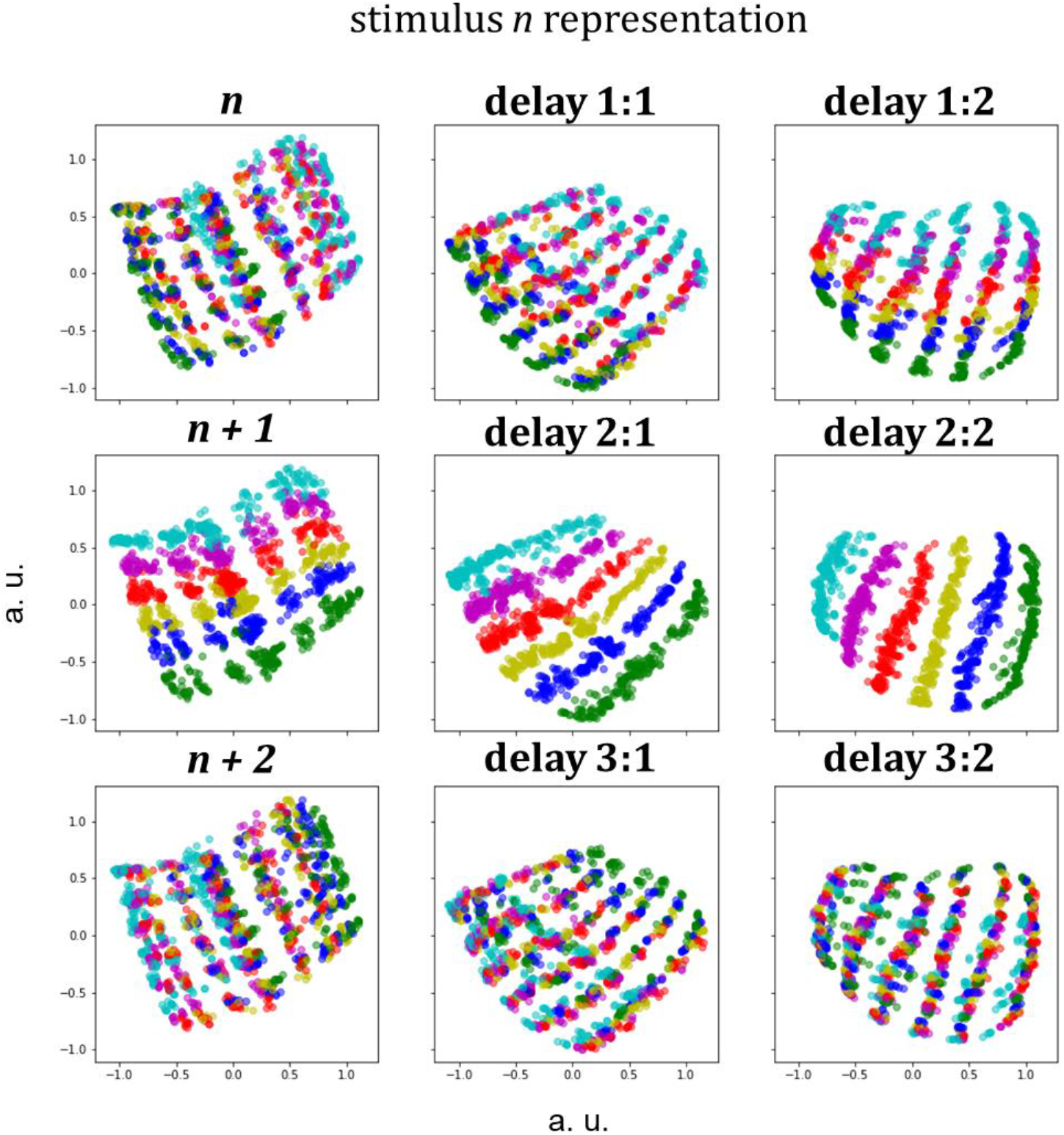
Example RNN trained with input following the basis function used to build IEMs in Wan et al. (2020). Shown is the 2D visualization of the LSTM hidden layer activity of this RNN. This is identical to simulation reported in the main text except that the inputs are not one-hot vectors; instead, they are specified by the IEM basis function: *R* = sin^6^(x) (e.g., for stimulus #3, input vector is [0.0156, 0.4219, 1, 0.4219, 0.0156, 0]). Results are qualitatively similar to RNNs reported in the main text (Figure 4).

**Figure S2.**
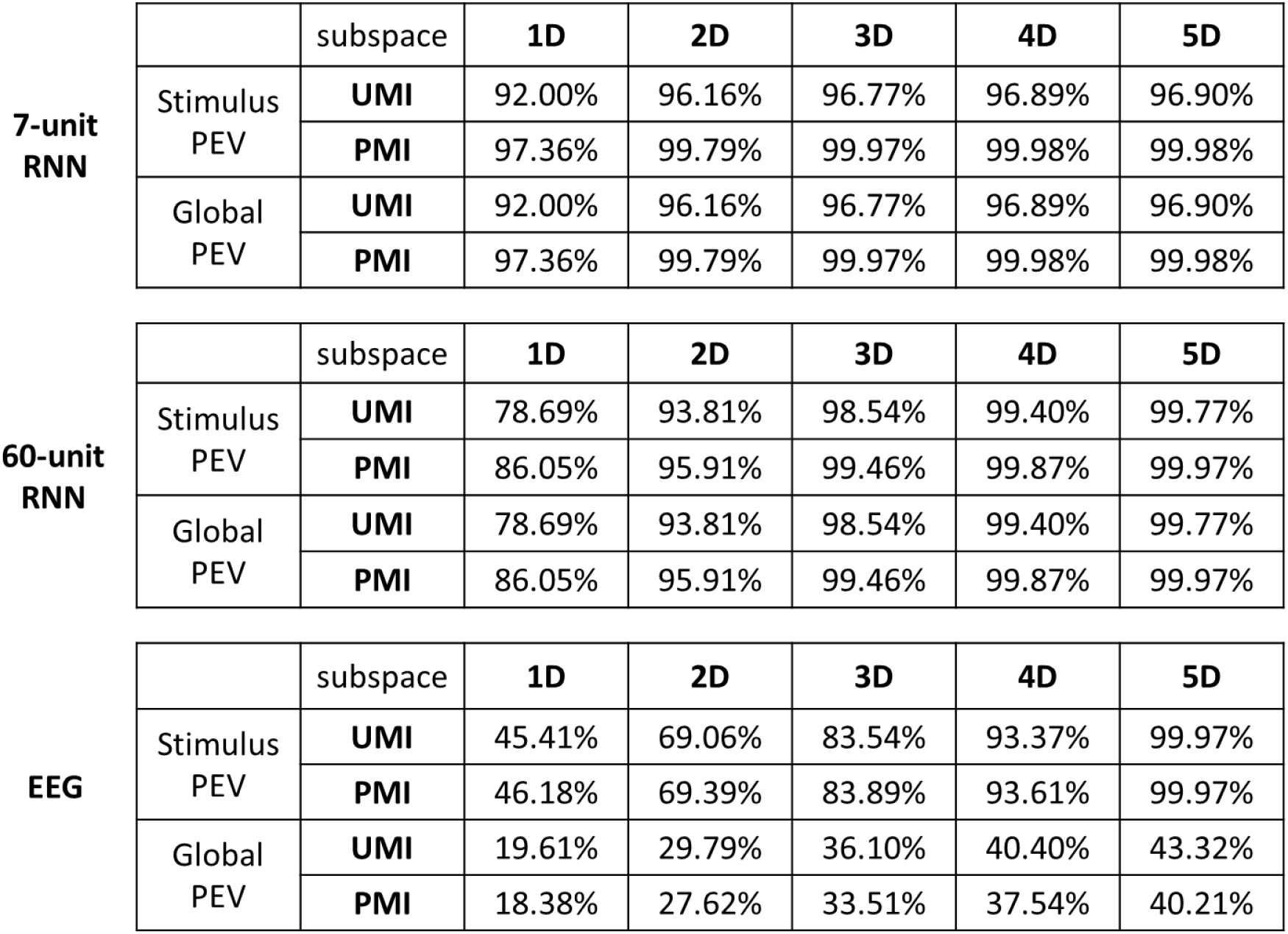
Cumulative percent variance explained (PEV) by top dPCs of the UMI and PMI subspaces for 7-unit RNN, 60-unit RNN and EEG data. The percentages of both stimulus and global variance explained are shown.

## References

Barak, O., & Tsodyks, M. (2014). Working models of working memory. Current Opinion in Neurobiology, 25, 20–24. https://doi.org/10.1016/j.conb.2013.10.008

Chatham, C. H., & Badre, D. (2015). Multiple gates on working memory. Current Opinion in Behavioral Sciences, 1, 23–31. https://doi.org/10.1016/j.cobeha.2014.08.001

Christophel, T. B., Iamshchinina, P., Yan, C., Allefeld, C., & Haynes, J.-D. (2018). Cortical specialization for attended versus unattended working memory. Nature Neuroscience, 21(4), 494–496. https://doi.org/10.1038/s41593-018-0094-4

Gardner, J. L., & Liu, T. (2019). Inverted encoding models reconstruct an arbitrary model response, not the stimulus. ENeuro, ENEURO.0363-18.2019. https://doi.org/10.1523/ENEURO.0363-18.2019

Gong, M., & Liu, T. (2020). Biased Neural Representation of Feature-Based Attention in the Human Frontoparietal Network. The Journal of Neuroscience, 40(43), 8386–8395. https://doi.org/10.1523/JNEUROSCI.0690-20.2020

Graves, A., Mohamed, A., & Hinton, G. (2013). Speech recognition with deep recurrent neural networks. 2013 IEEE International Conference on Acoustics, Speech and Signal Processing, 6645–6649. https://doi.org/10.1109/ICASSP.2013.6638947

Hochreiter, S., & Schmidhuber, J. (1997). Long Short-Term Memory. Neural Computation, 9(8), 1735–1780. https://doi.org/10.1162/neco.1997.9.8.1735

Kell, A. J., & McDermott, J. H. (2019). Deep neural network models of sensory systems: Windows onto the role of task constraints. Current Opinion in Neurobiology, 55, 121–132. https://doi.org/10.1016/j.conb.2019.02.003

Kingma, D. P., & Ba, J. (2017). Adam: A Method for Stochastic Optimization. ArXiv:1412.6980 [Cs]. http://arxiv.org/abs/1412.6980

Kobak, D., Brendel, W., Constantinidis, C., Feierstein, C. E., Kepecs, A., Mainen, Z. F., Qi, X.-L., Romo, R., Uchida, N., & Machens, C. K. (2016). Demixed principal component analysis of neural population data. ELife, 5, e10989. https://doi.org/10.7554/eLife.10989

Kruijne, W., Bohte, S. M., Roelfsema, P. R., & Olivers, C. N. L. (2020). Flexible Working Memory Through Selective Gating and Attentional Tagging. Neural Computation, 33(1), 1–40. https://doi.org/10.1162/neco_a_01339

LaRocque, J. J., Lewis-Peacock, J. A., Drysdale, A. T., Oberauer, K., & Postle, B. R. (2012). Decoding Attended Information in Short-term Memory: An EEG Study. Journal of Cognitive Neuroscience, 25(1), 127–142. https://doi.org/10.1162/jocn_a_00305

Lewis-Peacock, J. A., Drysdale, A. T., Oberauer, K., & Postle, B. R. (2011). Neural Evidence for a Distinction between Short-term Memory and the Focus of Attention. Journal of Cognitive Neuroscience, 24(1), 61–79. https://doi.org/10.1162/jocn_a_00140

Liu, T., Cable, D., & Gardner, J. L. (2018). Inverted Encoding Models of Human Population Response Conflate Noise and Neural Tuning Width. Journal of Neuroscience, 38(2), 398–408. https://doi.org/10.1523/JNEUROSCI.2453-17.2017

Lorenc, E. S., Vandenbroucke, A. R. E., Nee, D. E., de Lange, F. P., & D’Esposito, M. (2020). Dissociable neural mechanisms underlie currently-relevant, future-relevant, and discarded working memory representations. Scientific Reports, 10(1), 11195. https://doi.org/10.1038/s41598-020-67634-x

Manohar, S. G., Zokaei, N., Fallon, S. J., Vogels, T. P., & Husain, M. (2019). Neural mechanisms of attending to items in working memory. Neuroscience & Biobehavioral Reviews, 101, 1–12. https://doi.org/10.1016/j.neubiorev.2019.03.017

Mante, V., Sussillo, D., Shenoy, K. V., & Newsome, W. T. (2013). Context-dependent computation by recurrent dynamics in prefrontal cortex. Nature, 503(7474), 78–84. https://doi.org/10.1038/nature12742

Masse, N. Y., Yang, G. R., Song, H. F., Wang, X.-J., & Freedman, D. J. (2019). Circuit mechanisms for the maintenance and manipulation of information in working memory. Nature Neuroscience, 22(7), 1159. https://doi.org/10.1038/s41593-019-0414-3

Merrikhi, Y., Clark, K., Albarran, E., Parsa, M., Zirnsak, M., Moore, T., & Noudoost, B. (2017). Spatial working memory alters the efficacy of input to visual cortex. Nature Communications, 8(1), 15041. https://doi.org/10.1038/ncomms15041

Myers, N. E., Stokes, M. G., & Nobre, A. C. (2017). Prioritizing Information during Working Memory: Beyond Sustained Internal Attention. Trends in Cognitive Sciences, 21(6), 449–461. https://doi.org/10.1016/j.tics.2017.03.010

O’Reilly, R. C., & Frank, M. J. (2006). Making Working Memory Work: A Computational Model of Learning in the Prefrontal Cortex and Basal Ganglia. Neural Computation, 18(2), 283–328. https://doi.org/10.1162/089976606775093909

Panichello, M. F., & Buschman, T. J. (2021). Shared mechanisms underlie the control of working memory and attention. Nature. https://doi.org/10.1038/s41586-021-03390-w

Parthasarathy, A., Herikstad, R., Bong, J. H., Medina, F. S., Libedinsky, C., & Yen, S.-C. (2017). Mixed selectivity morphs population codes in prefrontal cortex. Nature Neuroscience, 20(12), 1770. https://doi.org/10.1038/s41593-017-0003-2

Richards, B. A., Lillicrap, T. P., Beaudoin, P., Bengio, Y., Bogacz, R., Christensen, A., Clopath, C., Costa, R. P., Berker, A. de, Ganguli, S., Gillon, C. J., Hafner, D., Kepecs, A., Kriegeskorte, N., Latham, P., Lindsay, G. W., Miller, K. D., Naud, R., Pack, C. C., … Kording, K. P. (2019). A deep learning framework for neuroscience. Nature Neuroscience, 22(11), 1761–1770. https://doi.org/10.1038/s41593-019-0520-2

Rose, N. S., LaRocque, J. J., Riggall, A. C., Gosseries, O., Starrett, M. J., Meyering, E. E., & Postle, B. R. (2016). Reactivation of latent working memories with transcranial magnetic stimulation. Science, 354(6316), 1136–1139. https://doi.org/10.1126/science.aah7011

Schneegans, S., & Bays, P. M. (2017). Restoration of fMRI Decodability Does Not Imply Latent Working Memory States. Journal of Cognitive Neuroscience, 29(12), 1977–1994. https://doi.org/10.1162/jocn_a_01180

Sprague, T. C., Adam, K. C. S., Foster, J. J., Rahmati, M., Sutterer, D. W., & Vo, V. A. (2018). Inverted Encoding Models Assay Population-Level Stimulus Representations, Not Single-Unit Neural Tuning. ENeuro, 5(3), ENEURO.0098-18.2018. https://doi.org/10.1523/ENEURO.0098-18.2018

Sprague, T. C., Boynton, G. M., & Serences, J. T. (2019). The Importance of Considering Model Choices When Interpreting Results in Computational Neuroimaging. ENeuro, 6(6). https://doi.org/10.1523/ENEURO.0196-19.2019

Sprague, T. C., Ester, E. F., & Serences, J. T. (2016). Restoring Latent Visual Working Memory Representations in Human Cortex. Neuron, 91(3), 694–707. https://doi.org/10.1016/j.neuron.2016.07.006

Stokes, M. G. (2015). ‘Activity-silent’ working memory in prefrontal cortex: A dynamic coding framework. Trends in Cognitive Sciences, 19(7), 394–405. https://doi.org/10.1016/j.tics.2015.05.004

Stokes, M. G., Muhle-Karbe, P. S., & Myers, N. E. (2020). Theoretical distinction between functional states in working memory and their corresponding neural states. Visual Cognition, 28(5-8), 420–432. https://doi.org/10.1080/13506285.2020.1825141

Sussillo, D., Churchland, M. M., Kaufman, M. T., & Shenoy, K. V. (2015). A neural network that finds a naturalistic solution for the production of muscle activity. Nature Neuroscience, 18(7), 1025–1033. https://doi.org/10.1038/nn.4042

Sutskever, I., Vinyals, O., & Le, Q. V. (2014). Sequence to Sequence Learning with Neural Networks. ArXiv:1409.3215 [Cs]. http://arxiv.org/abs/1409.3215

van Loon, A. M., Olmos-Solis, K., Fahrenfort, J. J., & Olivers, C. N. (2018). Current and future goals are represented in opposite patterns in object-selective cortex. ELife, 7, e38677. https://doi.org/10.7554/eLife.38677

Wan, Q., Cai, Y., Samaha, J., & Postle, B. R. (2020). Tracking stimulus representation across a 2-back visual working memory task. Royal Society Open Science, 7(8), 190228. https://doi.org/10.1098/rsos.190228

Yang, G. R., Joglekar, M. R., Song, H. F., Newsome, W. T., & Wang, X.-J. (2019). Task representations in neural networks trained to perform many cognitive tasks. Nature Neuroscience, 22(2), 297–306. https://doi.org/10.1038/s41593-018-0310-2

Yu, Q., Teng, C., & Postle, B. R. (2020). Different states of priority recruit different neural representations in visual working memory. PLOS Biology, 18(6), e3000769. https://doi.org/10.1371/journal.pbio.3000769

